# The evolution of social life in family groups

**DOI:** 10.1101/221192

**Authors:** Jos Kramer, Joël Meunier

## Abstract

Family life forms an integral part of the life-history of species across the animal kingdom, and plays a crucial role in the evolution of animal sociality. Our current understanding of family life, however, is almost exclusively based on studies that (i) focus on parental care and associated family interactions (such as those arising from sibling rivalry and parent-offspring conflict), and (ii) investigate these phenomena in the advanced family systems of mammals, birds, and eusocial insects. Here, we argue that these historical biases have fostered the neglect of key processes shaping social life in ancestral family systems, and thus profoundly hamper our understanding of the (early) evolution of family life. Based on a comprehensive survey of the literature, we first illustrate that the strong focus on parental care in advanced social systems has deflected scrutiny of other important social processes such as sibling cooperation, parent-offspring competition and offspring assistance. We then show that accounting for these neglected processes – and their changing role in the course of evolution – could profoundly change our understanding of the evolutionary origin and subsequent consolidation of family life. Finally, we outline how this diachronic perspective on the evolution of family living could provide novel insights into general processes driving social evolution. Overall, we infer that the explicit consideration of thus far neglected facets of family life, together with their study across the whole diversity of family systems, are crucial to advance our understanding of the processes that shape the evolution of social life.

## I. INTRODUCTION

Social life in family groups is a highly variable phenomenon that occurs widespread across the animal kingdom. Family groups can not only be found in vertebrates such as mammals, birds and (non-avian) reptiles, but also in numerous invertebrates including arthropods, molluscs and annelids (Clutton-Brock, 1991; Trumbo, 2012; Wong, Meunier, & Kölliker, 2013). Both within and across these taxa, families can vary tremendously in terms of composition, persistence and intimacy of social interactions (Klug, Alonso, & Bonsall, 2012; Trumbo, 2012). For instance, family groups can be composed of offspring and either their mother, their father, or both parents; they can last from only few hours to an entire lifetime; and they can range from temporary and facultative aggregations over cooperatively breeding groups to highly integrated eusocial societies featuring reproductive division of labour (Hölldobler & Wilson, 1990; Costa, 2006; Koenig & Dickinson, 2016).

The emergence of family life is commonly thought to constitute a transition from solitary to social life, and marks the initial step in the *major evolutionary transition* to eusociality (Maynard Smith & Szathmáry, 1995; Bourke, 2011). This is because the origin of family life entails the emergence of a novel – social – environment (cf. Badyaev & Uller, 2009; Uller, 2012) that can not only become an integral part of an organism’s life-history (Clutton-Brock, 1991; Gross & Clutton-Brock, 2005; Wong et al., 2013), but may also create long-lasting bonds between parents and their offspring. Such bonds preceded the evolution of many derived social behaviours (Darwin, 1871; Wilson, 1975; Royle, Smiseth, & Kölliker, 2012a), and thus likely drove the transformation of simple family systems to advanced animal societies. Eusocial societies, for instance, likely arose from family units in which offspring delayed dispersal and independent reproduction, and instead assisted their parents in raising younger siblings (Boomsma & Gawne, 2017). Studying family life can thus help elucidating factors that shape the evolution of complex animal societies (e.g. Wheeler, 1928; Michener, 1969; Wilson, 1975; Bourke, 2011), and more generally shed light on mechanisms that commonly promote the emergence and maintenance of social life in nature.

Despite its crucial role in social evolution, the origin and maintenance of family life is somewhat surprisingly often only touched upon indirectly in studies focusing on parental care (but see, for instance, (Falk et al., 2014; Jarrett et al., 2017). Parental care comprises a variety of traits ranging from gamete provisioning over nest construction to brood attendance and food provisioning (reviewed in Clutton-Brock, 1991; Costa, 2006; Smiseth, Kölliker, & Royle, 2012; Wong et al., 2013), and generally encompasses “any parental trait that enhances the fitness of a parent’s offspring, and that is likely to have originated and/or to be currently maintained for this function” (Smiseth et al., 2012). The expression of parental care often has a large impact on the fitness of both parents and offspring. In particular, parental care is beneficial to offspring, because it increases their quality and/or survival by neutralizing environmental hazards (Alonso-Alvarez & Velando, 2012; Klug & Bonsall, 2014). By contrast, parental care is often costly to parents, because it reduces their condition and/or survival (for instance as the result of an increased energy loss or elevated risk of predation), and thus ultimately diminishes their lifetime reproductive success (Trivers, 1972; Alonso-Alvarez & Velando, 2012). The far-reaching consequences associated with the expression of parental care make it the core feature of family life. Shedding light onto the circumstances that allow family members to gain sufficient (indirect) benefits to offset the costs of care (cf. Hamilton, 1964; Smiseth et al., 2012) has thus long been considered central in the study of social life in family groups (Clutton-Brock, 1991; Gross & Clutton-Brock, 2005).

However, parental care is but one of many facets of family life, and only a fraction of the other facets has received close scrutiny thus far. For instance, it is well known that the expression of care can prompt evolutionary conflicts (cf. Parker, Royle, & Hartley, 2002; Royle, Hartley, & Parker, 2004) that become apparent (i) if one parent tries to reduce its parental investment at the other parent’s expense *(parental antagonism;* Trivers, 1972; Lessells, 2012; Parker et al., 2015); (ii) if offspring compete with each other for limited parental resources *(sibling rivalry;* Mock & Parker, 1997; Roulin & Dreiss, 2012); and (iii) if offspring demand more care than the parents are willing to provide *(parent-offspring conflict;* Trivers, 1974; Kilner & Hinde, 2012; Kölliker et al., 2015). By contrast, processes such as sibling cooperation and parent-offspring competition only recently started to attract attention (e.g. Dreiss, Lahlah, & Roulin, 2010; Yip & Rayor, 2013; Falk et al., 2014; Schrader, Jarrett, & Kilner, 2015a; Kramer et al., 2017). This disparity arguably results from a strong bias toward studying family interactions in the derived social systems of birds and mammals. In these groups, young offspring are completely dependent on parental resources, and the substantial fitness effects of parental care that parallel this dependency typically prompt intense conflicts over the allocation of care (Clutton-Brock, 1991; Gross & Clutton-Brock, 2005). Derived family systems, however, only represent a small fraction of the diversity of family life in nature. Their predominance in studies of family interactions thus promotes the neglect of mechanisms that could play a greater role in less derived family systems. Moreover, the central role of parental care arguably deflects scrutiny of fitness effects that are typically masked by the benefits and costs of (conflicts over the allocation of) parental care. The strong focus on parental care and its expression in altricial species hence likely distorts our understanding of the evolutionary drivers of the emergence and consolidation of family life, and could ultimately obscure their role in the (early) evolution of animal sociality.

Here, we advocate the direct study of family life as an integrative approach to elucidating the role of parental behaviours and other family interactions in the evolution of animal sociality. To this end, we (i) illustrate the downsides of a narrow focus on parental care by reviewing how thus far neglected types of family interaction can shape the cost-benefit ratio of family life. We then (ii) outline how accounting for these overlooked mechanisms – and their changing role in the course of evolution – could improve our understanding of the evolutionary origin and consolidation of family life. Finally, (iii) we discuss how this diachronic perspective on the evolution of family living could provide general insights into the mechanisms driving social evolution. Understanding the evolution of family life requires a complete picture of all factors that affect its fitness consequences across taxonomical groups. Albeit doubtlessly very important, parental care and its repercussions in advanced family systems only cover part of the canvas.

## II. THE SEMANTICS OF FAMILY LIFE

Somewhat surprisingly, there is no strict consensus among behavioural ecologists as to what constitutes a family. In studies on cooperative breeding, the term family is typically restricted to cases where mature offspring forgo dispersal and independent reproduction, and instead continue to interact regularly with their parents (Emlen, 1994, 1995; Covas & Griesser, 2007; Drobniak et al., 2015). This narrow definition helps to identify transitional stages in the evolution of cooperative breeding (a form of family-living characterized by reproductive cooperation; Drobniak et al., 2015). Yet, this definition also excludes the vast diversity of (less enduring) associations between parents and their *immature* offspring. A broader meaning of the term «family» is thus frequently implied in studies on parental care (cf. Clutton-Brock, 1991; Gross & Clutton-Brock, 2005; Schrader et al., 2015b; Duarte et al., 2016; Jarrett et al., 2017). Here, we formalize this view by defining a family as “an association of one or both caring parent(s) with their offspring”. This broad definition of the term family closely matches its colloquial meaning, and allows us to outline a general perspective that covers all types of (non-random) parent-offspring association. We suggest using more narrowly define terms such as *nuclear family* and *extended family* to delineate families of a particular composition. Specifically, we propose to use the term *nuclear family* to delineate the vast majority of family systems that consist of one or both caring parent(s) and offspring of a single reproductive attempt. Conversely, we suggest using the term *extended family* to delineate families consisting of a nuclear family and their close relatives, such that the extended family also comprises grandparents, siblings of the parents, and/or offspring of at least one additional reproductive attempt. Many societies of cooperatively breeding birds, mammals, and eusocial insects are examples of such extended families.

Family systems may not only differ in terms of composition, but also in terms of the extent to which parental care is integrated into offspring development. This latter difference is broadly captured by the classification into species with altricial and precocial young (from now on referred to as altricial and precocial species, respectively). In altricial species, the phenotypic integration of parental care is advanced to such an extent that juveniles cannot survive without receiving at least some care early during their life. Family life in altricial species is therefore obligatory (Clutton-Brock, 1991). Prime examples of such altricial species are found among mammals, passerine birds, and eusocial insects. In precocial species, on the other hand, this phenotypic integration is limited, and offspring can survive in the absence of – nonetheless beneficial – care due to an early development of their capability to forage independently. Family life in precocial species is therefore facultative (Smiseth, Darwell, & Moore, 2003; Kölliker, 2007). Ducks, plovers, and quails, as well as many subsocial insects (such as burying beetles and earwigs) feature precocial young. Interestingly, the altricial-precocial spectrum broadly coincides with the classification into evolutionarily derived vs. non-derived family systems. In particular, altricial family systems are always derived (and derived systems typically altricial), since the high phenotypic integration of parental care characteristic of such systems only arises *after* the emergence of family life (section III.3; Kölliker, 2007; Uller, 2012). Conversely, precocial species are typically less derived (and non-derived systems are always precocial), since they feature a lower degree of phenotypic integration, and thus more closely resemble an ancestral state during which offspring were (still) largely independent of parental care (Smiseth et al., 2003; Kölliker, 2007). We are aware that these dichotomic classifications only draw a rough picture of the diversity of family systems. We nevertheless retain them here, because their generality makes them useful in our discussion of the general trends shaping the evolution of family life.

## III. THE NEGLECTED FACETS OF FAMILY LIFE

Family living is a form of group living. The various fitness effects inherent to all types of group-living – such as the costs of increased competition and the benefits of cooperative foraging -should therefore also occur during family life (Alexander 1974; Krause and Ruxton 2002). However, instead of investigating the full range of possible cooperative and competitive family interactions, the last decades of research on family life predominantly focused on the fitness effects of a single type of parent-offspring cooperation – parental care – as well as on the conflicts over (and the cooperation in) its allocation (see Figure 1; reviewed in Clutton-Brock, 1991; Royle, Smiseth, & Kölliker, 2012b). This implicit equalization of the fitness effects of parental care with the fitness effects of family life has led to the neglect of three potentially important dimensions of social interactions within the family: (i) sibling cooperation, (ii) parent-offspring competition, and (iii) offspring assistance (here defined as cooperative acts of offspring to the benefit of their parents). A notable exception to this general trend are studies on the highly derived, extended families of cooperative breeders, in which these mechanisms have been explored (but see section IV. 2). Below, we review examples of the neglected facets of family life, and highlight that their fitness effects are often concealed by the relatively greater effects of parental care. Accounting for the effects of these mechanisms is nevertheless crucial, since they could augment or diminish the benefits and costs of parental care, and thus tip the scales in favour (or in disfavour) of the emergence and consolidation of family life.

**Figure 1.**
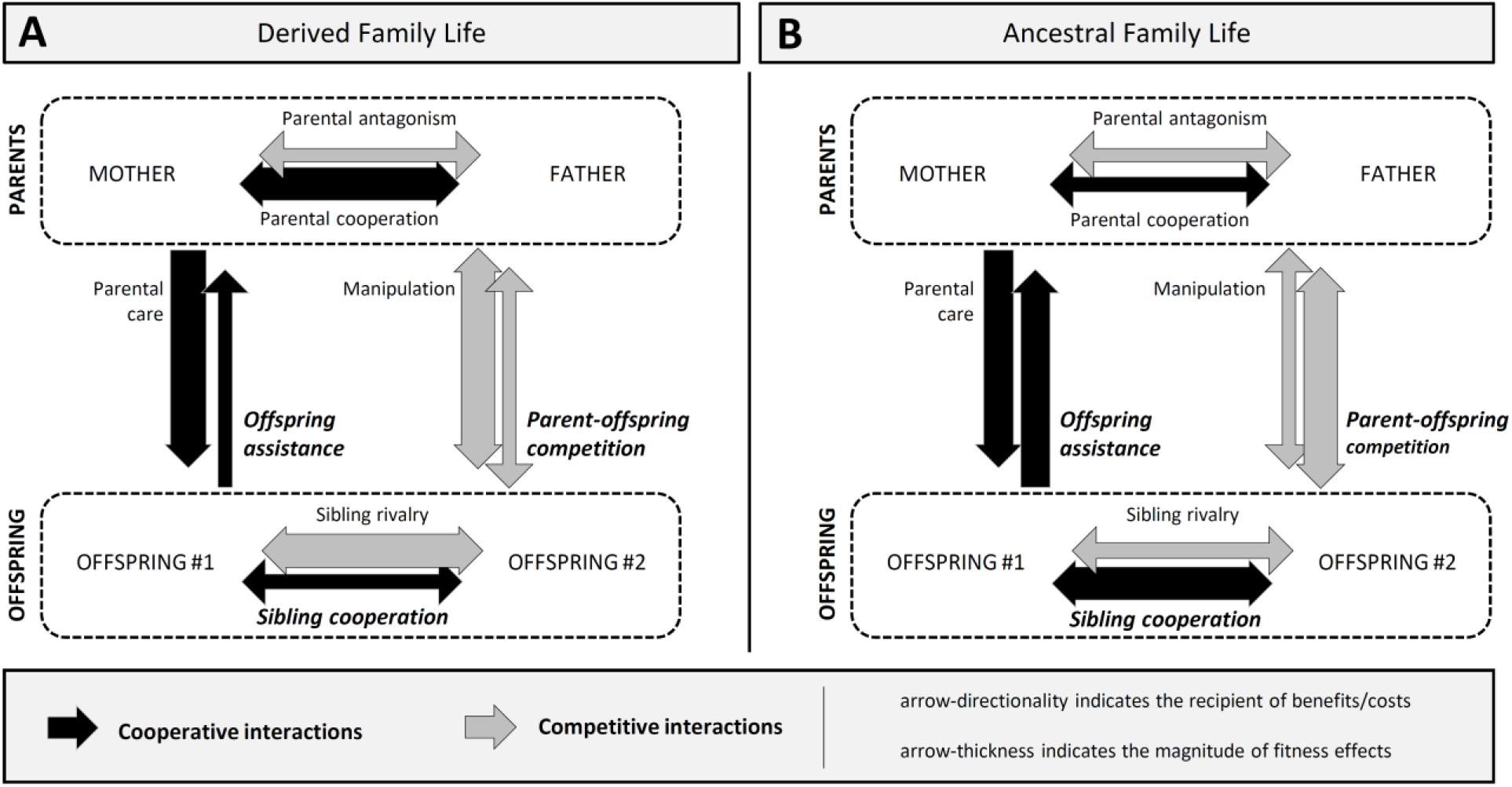
Social interactions during family life. Depicted are cooperative and competitive interactions (represented, respectively, by black and grey arrows) that can potentially occur among family members in (A) derived and (B) ancestral family systems. Research on family interactions has traditionally focused on altricial vertebrates and eusocial insects, and typically investigated the expression and fitness effects of parental care and the conflicts over (and cooperation in) its allocation. While this strong focus is understandable in the light of the often substantial fitness effects of these phenomena (indicated by the thickness of the corresponding arrows) in derived family systems, it has inadvertently fostered the neglect of other facets of family life (in bold italic print). However, these neglected facets might have played a crucial role in shaping ancestral forms of family life. Notably, the social dynamics in ancestral family systems might be very similar to the dynamics in the extended families of many cooperative breeders.

### (1) Sibling cooperation

Most studies of sibling interactions thus far investigated the conspicuous competitive behaviours of juvenile birds and mammals that compete over their access to limited parental resources (reviewed in Mock & Parker, 1997; Roulin & Dreiss, 2012). However, sibling interactions are not competitive by default, and an increasing number of studies report an unexpected diversity of cooperative interactions (or by-product mutualism) among altricial as well as precocial juveniles (reviewed in Roulin & Dreiss, 2012). Indeed, sibling cooperation is not only a hallmark of termite societies, where larvae, nymphs, workers and soldiers are all juveniles (Eggleton, 2011), but also occurs in the house wren *Troglodytes aedon*, where offspring postpone fledging to the benefit of their younger siblings (Bowers, Sakaluk, & Thompson, 2013); in the King Penguin *Aptenodytes patagonicus*, where huddling improves the juveniles’ thermoregulation (Barré, 1984); in the spotted hyaena *Crocuta crocuta*, where offspring form coalitions with litter-mates against unrelated juveniles (Smale et al., 1995); and in the Mississippi kite *Ictinia mississippiensis* and the ambrosia beetle *Xyleborinus saxesenii*, where offspring express mutual cleaning (‘allo-preening’; Botelho, Gennaro, & Arrowood, 1993; Biedermann & Taborsky, 2011).

Intriguingly, juveniles can also cooperate in resource acquisition. In altricial species, such cooperation typically aims at improving the juveniles’ access to parental provisioning (cf. Forbes, 2007). For instance, altricial juveniles sometimes refrain from interfering with their siblings’ feeding attempts (e.g. in the blue-footed booby *Sula nebouxii;* Anderson & Ricklefs, 1995) and may even offer parentally-provided food items to their siblings (e.g. in the barn owl *Tyto alba;* Marti, 1989). Moreover, they can coordinate their begging behaviour to increase the parents’ feeding rate (e.g. in the in the black-headed gull *Larus ridibundus* and the banded mongoose *Mungos mungo;* Johnstone, 2004; Mathevon & Charrier, 2004; Bell, 2007), or negotiate their share of parental resources to avoid the greater costs of unrestrained sibling rivalry (e.g. in the barn owl *T. alba*, the spotless starling *Sturnus unicolor* and the meerkat *Suricata suricatta;* Roulin, 2002; Johnstone & Roulin, 2003; Bulmer, Celis, & Gil, 2008; Madden et al., 2009; Dreiss et al., 2010). By contrast, cooperation in resource acquisition among precocial juveniles can – at least in principle – occur independently of parental provisioning. For instance, food sharing occurs even without parental involvement in the European earwig *Forficula auricularia* (Falk et al., 2014; Kramer, Thesing, & Meunier, 2015) and in many social spiders such as the huntsman spider *Delena cancerides* (Yip & Rayor, 2013, 2014).

In both altricial and precocial species, the fitness effects of sibling cooperation might often be concealed by the effects of parental care. In line with this assumption, it has recently been shown that larval mass in the burying beetle *Nicrophorus vespilloides* peaks at a higher larval density in the absence of care. This suggests that parental care usually masks the beneficial effect of initial increases in larval density on the brood’s ability to penetrate and use the breeding carcass (Schrader et al., 2015a). This notwithstanding, the diverse forms and broad taxonomical distribution of cooperative behaviours among juveniles suggest that sibling cooperation is not only important during the adult life stage (cf. Wilson, 1971; Clutton-Brock, 1991; Koenig & Dickinson, 2004), but might also play a crucial role in the evolution of family life (see section III).

### (2) Parent-offspring competition

Competition between parents and their offspring occurs whenever the consumption of an essential resource by the parents limits the availability of this resource to their offspring – or vice versa. This form of kin competition typically arises with the onset of offspring foraging, and has been predicted to promote offspring dispersal and thus the breakup of family units (Hamilton & May, 1977; Comins, Hamilton, & May, 1980; West, Pen, & Griffin, 2002). Direct evidence for these effects in altricial species is as yet scarce. However, experimental food removal in the western bluebird *Sialia mexicana* reduced the number of sons that remained with their parents until after winter, suggesting that parent-offspring competition can indeed promote the breakup of family units (Dickinson & McGowan, 2005). Similarly, food supplementation delayed offspring dispersal in the carrion crow *Corvus corone corone*, indicating that parent-offspring competition could select against the evolution of cooperative breeding (Baglione et al., 2006). In further support of this notion, resource depletion during the breeding season likely leads to competition between breeders and helpers in the chestnut-crowned babbler *Pomatostomus ruficeps*, and thus overall increases the costs of group-living (Sorato, Griffith, & Russell, 2016).

In analogy to its effects in altricial species, parent-offspring competition can also affect offspring dispersal and the duration of family life in precocial species. For instance, parent-offspring competition has been shown to promote offspring dispersal in the solitary common lizard *Lacerta vivipara* (Léna et al., 1998; Le Galliard, Ferrière, & Clobert, 2003; Cote, Clobert, & Fitze, 2007). Conversely, the prolonged presence of fathers has been shown to reduce offspring survival under food limitation in *N. vespilloides*, a burying beetle with biparental care in which both parents feed on the breeding carcass (Scott & Gladstein 1993; Boncoraglio & Kilner 2012). This latter finding suggests that father-offspring competition might offset the benefits of paternal care, and thus offers a potential explanation as to why fathers typically leave the brood earlier than mothers in this species. Intriguingly, parent-offspring competition in precocial species might even entirely negate the benefits of family living under certain harsh conditions. In line with this hypothesis, mother-offspring competition under food limitation has been shown to render maternal presence detrimental to offspring survival in uniparental families of the European earwig *F. auricularia* (Meunier & Kölliker, 2012; Kramer et al., 2017).

Particularly in precocial species, such costs of parent-offspring competition are likely often concealed by the benefits of parental care. Indeed, carcasses guarded by *N. vespilloides* fathers are less likely to be taken over by conspecifics, suggesting that the costs of father-offspring competition are typically offset by the benefits of offspring defence against infanticide through conspecifics (Scott & Gladstein, 1993). Similar benefits of parental care might also explain why *F. auricularia* offspring do not disperse earlier under resource limitation (Wong & Kölliker, 2012). Finally, note that such masking effect of parental care are likely less pronounced in altricial species, since the benefits of parental care often decrease towards the end of family life (cf. Bateson, 1994), and will thus often be limited once parent-offspring competition arises. Given its early onset and multifaceted role in precocial species, parent-offspring competition might play a crucial role in the evolution of family life (see section III).

### (3) Offspring assistance

Cooperation between parents and their offspring is prominently featured in a plethora of studies on parental care (reviewed in Clutton-Brock, 1991; Royle et al., 2012b). However, parent-offspring cooperation is not a one-way road and can also involve cooperative behaviours (or by-product mutualism) that offspring direct towards their parents. Such *offspring assistance* is pervasive in the extended families of cooperative breeders, where adult offspring often assist their parents in raising younger siblings (Wilson, 1971; Bourke & Franks, 1995; Cockburn, 1998; Koenig & Dickinson, 2016). Yet offspring assistance during family life can also be performed by juveniles. Among altricial species, it frequently occurs in eusocial insects where larvae/nymphs can fulfil crucial roles for colony functioning (reviewed in Eggleton, 2011; Schultner, Oettler, & Helanterä, 2017), for instance by defending the colony – and thus the reproductives – as soldiers (in virtually all termites; (Howard & Thorne, 2011); by taking over gallery extension and the compressing of waste into compact balls (in the ambrosia beetle *X. saxesenii;* Biedermann & Taborsky, 2011); or by acting as “communal stomach” (cf. Wheeler, 1918; Dussutour & Simpson, 2009) that provisions the queen with secretions necessary for protein degradation (in the metricus paper wasp *Polistes metricus;* Hunt, 1984) or sustained egg production (in pharaoh ant *Monomorium pharaonis;* Børgesen, 1989; Børgesen & Jensen, 1995).

Apart from its role in altricial species with highly complex societal organization, the notion of offspring assistance has received little attention. However, recent findings indicate that parents can also benefit from offspring assistance in precocial species. For instance, parents might benefit from their offspring’s investment into shared immune traits (*social immunity;* Cremer, Armitage, & Schmid-Hempel, 2007; Meunier, 2015), independent foraging, or defence against predation (Krause & Ruxton, 2002). In line with the former notion, faeces of caring mothers exhibit a lower antifungal activity than those of non-caring females in the European earwig *F. auricularia*, suggesting that mothers might downregulate or at least not compensate for the reduction in their own investment into nest sanitation during family life, and instead rely on the superior antifungal properties of the faeces of their juveniles (Diehl et al., 2015). Conversely, delayed juvenile dispersal improves the survival of tending mothers in the subsocial spider *Anelosimus studiosus* (Jones & Parker, 2002), a finding that might reflect benefits of offspring investment into prey capture or into the maintenance of the communal web. Indeed, offspring assist in web construction in many social spiders (Yip & Rayor, 2014), suggesting that mothers could regularly benefit by reducing their own investment. Although such benefits of offspring assistance for parents might be concealed by the costs of parental care, they could nevertheless have a significant impact on the evolution of family interactions and, more generally, on the emergence of social life in family units (see section III). Investigating the role of offspring assistance and the other neglected facets of family life is thus crucial to advance our understanding of the evolution of social life in family groups.

## III. THE (EARLY) EVOLUTION OF FAMILY LIFE

The evolution of family life generally presumes that the fitness benefits of family living outweigh the costs of a prolonged association of the family members (Alexander, 1974; Clutton-Brock, 1991; Klug et al., 2012). However, the impact of the processes mediating these benefits can change over evolutionary time (Smiseth et al., 2003; Falk et al., 2014; Royle, Alonzo, & Moore, 2016). This is because the current benefits associated with a trait (such as a parental behaviour) do not necessarily reflect the adaptive value of this trait in an ancestral state (Williams, 1966). For instance, the high benefits associated with parental food provisioning in derived family systems typically reflect the dependency of offspring on food provided by the parents, a state that only evolved *after* the emergence of parental provisioning. The benefits of parental provisioning are thus likely less pronounced in non-derived family systems (Smiseth et al., 2003; Klug et al., 2012; Royle et al., 2012a). Conversely, mechanisms playing a limited role in derived systems might have a more prominent role in less derived systems (section III.2.b). Understanding the evolution of family living therefore requires a complete picture of the mechanisms promoting family life in both derived and non-derived family systems.

However, instead of investigating the full range of mechanisms across different family systems, the last decades of empirical research mostly focused on investigating the current benefits and costs of parental care in the derived family systems of altricial vertebrates (reviewed in Clutton-Brock, 1991; Royle et al., 2012b). By contrast, the fitness effects of family interactions in precocial species, which feature facultative forms of family life reminiscent of an ancestral state, have received comparably little attention (but see, for instance, Eggert, Reinking, & Müller, 1998; Zink, 2003; Salomon, Schneider, & Lubin, 2005; Kölliker, 2007). Similarly, theoretical approaches have thus far only indirectly explored the evolution of family life, since they typically investigated the influence of life-history characteristics, co-evolutionary dynamics, or environmental conditions on the evolutionary origin and maintenance of parental care (Wilson, 1975; Tallamy, 1984; Tallamy & Wood, 1986; Bonsall & Klug, 2011a; Klug et al., 2012). As a corollary of this narrow focus on parental care, our current understanding of the early evolution of family life remains fragmentary. In the following section, we address this fundamental issue. Specifically, we review the factors promoting the emergence and subsequent consolidation of family life, and demonstrate that integrating the costs and benefits of thus far overlooked facets of family life in particular – and the study of precocial family systems in general – could entail major changes in our understanding of the evolution of family life.

## (1) The emergence of family life

### (a) The standard account: the evolution of post-hatching parental care

The evolutionary emergence of family life has typically been explored indirectly in studies endeavouring to understand which factors favoured the extension of pre-hatching parental care beyond the time of offspring emergence (e.g. Lack 1968; Clutton-Brock 1991; Smiseth et al. 2012). These studies suggest that the emergence of parental care – and thus family life – requires the concurrence of factors that jointly make sustained social interactions among family members possible and – should the occasion arise – able to spread in the population (Klug et al., 2012). The initial step in the emergence of family life is promoted by life-history characteristics ensuring that social behaviours are primarily directed toward family members (Tallamy & Wood, 1986; Lion & van Baalen, 2007). This propensity to mainly interact with family members increases the scope for the evolution of cooperative behaviours (such as parental care and sibling cooperation) by reducing the likelihood that such behaviours are misdirected toward non-kin (Hamilton, 1964; Lion & van Baalen, 2007). Hence, family life is most likely to emerge if parents and offspring recognize each other (e.g. by means of kin or familiarity recognition; cf. Evans 1998; Fellowes 1998; Dobler and Kölliker 2011) or if they frequently encounter each other (e.g. due to limited dispersal; Hamilton 1964a; Lion and van Baalen 2007). Additionally, the emergence of family life can be promoted by the presence of precursors of post-hatching care (Tallamy & Wood, 1986; Royle et al., 2012a). In line with this idea, the evolution of offspring attendance and guarding in cooperative breeders has been suggested to derive from ancestral defensive or aggressive behaviours (Tallamy, 1984). Similarly, parental provisioning during family life might have evolved via selection acting on – and modifying – self-feeding behaviours (Cunningham et al., 2016), and some effector molecules in social immunity might have been recruited from a function in personal immunity (Palmer et al., 2016).

Once the preconditions for the emergence of family life are met, effects of (additional) life-history characteristics and environmental conditions jointly determine whether it can spread in the population against the background of the prevalent solitary lifestyle (Tallamy, 1984; Clutton-Brock, 1991; Klug et al., 2012). In particular, environmental conditions – including the spatial and temporal availability of limited resources and the presence of predators or parasites (reviewed in Wilson, 1975; Krause & Ruxton, 2002; Botterill-James et al., 2016; see also Botterill-James et al., 2016) – typically modify the impact of basic life-history conditions (such as stage-specific mortality and maturation rates) on the benefits and costs of family interactions (Bonsall & Klug, 2011a; Klug et al., 2012). For instance, harsh conditions and the concomitant intense competition for limited resources have been predicted to increase the mortality rate of solitary individuals (Wilson, 1975; Clutton-Brock, 1991). This, in turn, should promote the evolution of parental care and thus family life, because the uncertain prospects of future reproduction decrease the relative costs of care to adults (Klug & Bonsall, 2010; Bonsall & Klug, 2011a), and increase its potential benefits to offspring (Webb et al., 2002; Klug & Bonsall, 2010). However, empirical findings are sometimes at odds with these predictions. For instance, harsh conditions negate rather than increase the usual benefits of maternal presence and thus family life in the European earwig *F. auricularia* (Meunier & Kölliker, 2012; Kramer et al., 2017). The limited predictive power of the standard account of the evolution of family life (cf. Costa, 2006; Trumbo, 2012; Capodeanu-Nägler et al., 2016) might partly reflect that environmental conditions, life-history characteristics, and the benefits and costs of parental care often interact in unexpected ways (Bonsall & Klug, 2011a, 2011b; Meunier & Kölliker, 2012). However, we believe that it also reflects an excessive focus on a subset of family interactions, and their expression in a subset of family systems.

### (b) An extended account: the role of the neglected facets of family life

The standard account for the evolutionary origin of family life solely focuses on the extension of parental care beyond offspring emergence, and thus inadvertently neglects the role of other social interactions within the nascent family. However, these neglected facets could have a profound influence on family life. In particular, parent-offspring competition (and its potential knock-on effects on sibling rivalry and parental antagonism) could impede the evolution of family life by reducing the potential benefits of care (Meunier & Kölliker, 2012; Kramer et al., 2017). Conversely, both sibling cooperation and offspring assistance could promote the emergence of family living by, respectively, augmenting the (initially limited) benefits of care to offspring, and offsetting some of its costs to parents (cf. Falk et al., 2014; Kramer et al., 2015). For instance, sibling cooperation during foraging could promote reciprocal food sharing (such as in the vampire bat *Desmodus rotundus;* Wilkinson, 1984; Carter & Wilkinson, 2013), and thus provide a mechanism for insurance against variability (Koenig & Walters, 2015). Intriguingly, these forms of cooperation could themselves evolve from by-product benefits (such as predator dilution effects; Krause & Ruxton, 2002) arising in offspring aggregations.

The benefits of by-product mutualism or sibling cooperation in such offspring aggregations could also affect the initial duration of family life. In particular, they could offer an additional incentive (or even an alternative reason; see section IV.1) for offspring to delay dispersal from their natal site (Kramer et al., 2015), and might thus allow extended periods of family life right from the start. This scenario contrast with the standard account for the evolution of family life, where the “simple” extension of parental care beyond offspring emergence (cf. Michener, 1969; Costa, 2006) should initially only allow for brief periods of family life. This is because the standard account neglects the potential impact of cooperation among offspring, and thus implies that offspring in recently evolved family systems should (still) tend to disperse soon after hatching to avoid the impending competition with their siblings and parents (West et al., 2001, 2002). Longer periods of family-living would only arise secondarily, where the benefits of offspring attendance and other early forms of parental care select for delayed offspring dispersal. From an offspring’s point of view, family life is classically thought to evolve *despite of* the presence of competing siblings (cf. Mock & Parker, 1997; Roulin & Dreiss, 2012). However, the occurrence and potential role of sibling cooperation suggests that family life might rather emerge – or at least be initially favoured – *because of* the presence of siblings.

Similar to the fitness effects of early forms of parental care (see section III.1.a), the impact of other facets of family life likely depends on life-history characteristics and the prevailing environmental conditions. Costs of parent-offspring competition, for instance, will be greatest if parents and offspring feed on the same resources, and simultaneously forage in the same area. Conversely, the benefits of sibling cooperation might be greatest if offspring forage independently of each other, since this would decrease sibling rivalry (cf. Mock & Parker, 1997), and thus increase the incentive of juveniles to cooperate with each other (Frank, 1998, 2003). Finally, the possible spectrum of different types of sibling cooperation and offspring assistance is likely subject to developmental constraints (cf. Maynard Smith et al., 1985), where certain types of behaviours cannot be performed effectively by immatures. Besides these life-history traits, the (environmentally determined) availability of limited resources is likely a crucial factor shaping the fitness effects of the neglected facets of family life. This is because resource limitation would both increase the scope for parent-offspring competition, and decrease the propensity of juveniles to cooperate with their siblings or parents (West et al., 2002; Frank, 2003; see also section III.2.b). Such harsh conditions might thus hamper the evolution of family life despite the expected high benefits of parental care (Webb et al., 2002; Klug & Bonsall, 2010). Overall, such as yet poorly explored effects might help explaining why even closely related species exposed to ostensibly identical conditions often differ in the occurrence and nature of family interactions (cf. Costa, 2006; Trumbo, 2012; Capodeanu-Nägler et al., 2016).

## (2) The consolidation of family life

### (a) The standard account: the evolution of elaborate care

After the emergence of family units, coevolutionary feedback-loops between parental and offspring traits are expected to promote the evolution and diversification of parental care, and thus to lead to the rapid consolidation of family life (Wolf, Brodie III, & Moore, 1999; Kölliker, Royle, & Smiseth, 2012; Uller, 2012; Jarrett et al., 2017). For instance, the initial evolution of parental provisioning may trigger evolutionary changes in other components of care as well as in offspring traits, allowing parents to choose safer nest sites, but also increasing the competition among offspring for parentally provided food. This increased sibling rivalry may, in turn, further advance the evolution of parental provisioning, thereby closing the coevolutionary feedback-loop between parental provisioning, the choice of safer nest sites, and sibling rivalry (Smiseth, Lennox, & Moore, 2007; Gardner & Smiseth, 2011). Such mutual reinforcement between parental and offspring traits has been predicted to promote a unidirectional trend from simple ancestral forms toward complex forms of family life by fostering an increasingly tight phenotypic integration of parental care and offspring development (Wilson, 1975; Gardner & Smiseth, 2011; Kölliker et al., 2012; Uller, 2012; Royle et al., 2016). In the highly-derived family systems of altricial species, this phenotypic integration is advanced to such an extent that juveniles cannot survive without at least some care early in their life (Kölliker, 2007; Uller, 2012).

### (b) An extended account I: the (changing) role of the neglected facets of family life

The increasingly tight integration of parental care into offspring development that evolves during the consolidation of family life could have a profound effect on the relative importance of the neglected facets of family life. For instance, the evolution of parental provisioning and the concomitant increased reliance of offspring on parentally provided food likely leads to a delayed onset of offspring foraging (cf. Gardner & Smiseth, 2011), and should thus reduce the scope for competition between parents and their offspring. As a result, the impact of parent-offspring competition on family dynamics might steadily decline in the course of the consolidation of family life (Kramer et al., 2017). Similarly, siblings might be most likely to cooperate with each other as long as they are (still) largely independent of parental care. This is because an increased dependency on care is typically paralleled by increased sibling rivalry (Gardner & Smiseth, 2011), and should thus decrease the levels of sibling cooperation (Frank, 1998, 2003). Finally, an increased offspring dependency is likely also accompanied by greater developmental constraints (cf. Maynard Smith et al., 1985) on the type of social behaviours that the immature juveniles can perform, suggesting that both sibling cooperation and offspring assistance might occur less frequently in altricial than in precocial species. Overall, these considerations indicate that parent-offspring competition, sibling cooperation, and offspring assistance might fulfil crucial roles in ancestral family systems, but could lose ground where the consolidation of family life promotes an increasingly tight phenotypic integration of parental care into offspring development.

While the role of parent-offspring competition, sibling cooperation, and offspring assistance in nuclear families thus far received little attention, their impact on the evolution of the extended families of cooperative breeders has been more thoroughly explored (e.g. Bourke & Franks, 1995; Baglione et al., 2006; Koenig & Dickinson, 2016; Sorato et al., 2016). Interestingly, all three facets play a prominent role in shaping these systems: parent-offspring competition can impede the evolution of cooperative breeding (Baglione et al., 2006; Sorato et al., 2016), siblings within breeding groups frequently cooperate with each other (e.g. during group foraging or in the defence against predation), and offspring assistance in the form of alloparental care (often called ‘help’) is the very foundation of cooperative breeding (Skutch, 1935; Cockburn, 1998; Koenig & Dickinson, 2016). However, while these mechanisms usually involve juveniles in nuclear families, they typically involve adult offspring in cooperative breeders. Notably, the resurgence of these mechanisms in cooperative breeders after their demise during the consolidation of (nuclear) family life is in line with a key role of offspring dependency in determining their occurrence. Like juveniles in ancestral family systems, adult offspring in cooperative breeders are largely independent of parental care, a situation that not only promotes parent-offspring competition, but also shifts competition towards a global(population-wide) scale (cf. West et al., 2002), and thus prevents the high levels of sibling rivalry that usually reduce the likelihood that sibling cooperation and offspring assistance occur (see above). The putatively similar role of these mechanisms in ancestral and cooperatively breeding families suggests that the extensive literature on the evolution of cooperative breeding could inform studies of the emergence of family life from a solitary state. In particular, the distinction between a helper’s decision to stay and its subsequent decision to provide alloparental care (e.g. Ekman & Tegelström, 1994; Griesser et al., 2017) could be applied to the evolution of family units, and might then suggest that the initial formation of family units is not necessarily (only) driven by the benefits of parental care (see also section IV.1).

### (c) An extended account II: the rocky road to complex family systems

Far from being restricted to elucidating the role of the three neglected facets of family life, the study of family interactions in precocial species can also shed light on other aspects of the evolution of family living. For instance, it might help explaining why simple family life still abounds across taxa (e.g. Tallamy & Schaefer, 1997; Lin, Danforth, & Wood, 2004; Filippi et al., 2009) despite the expected trend towards complex family systems (Wilson, 1975; Gardner & Smiseth, 2011; Kölliker et al., 2012; Uller, 2012; Royle et al., 2016). In general terms, this mismatch between theoretical expectations and empirical findings indicates that some as yet unknown factors counteract the consolidation of family life and thus prevent an increase in social complexity. We recently showed that long-term and transgenerational costs of parental loss (such as an impaired development of juveniles) are not restricted to altricial family systems (e.g. Harlow & Suomi, 1971; Gonzalez et al., 2001; Fleming et al., 2002; Andres et al., 2013), but can also occur in precocial species (Thesing et al., 2015). This finding suggests that the mortality rate of parents *during* family life could be one of the factors counteracting its consolidation. This is because even though precocial juveniles can survive the early death of their parents, they will still suffer (non-lethal) consequences of parental loss. Accordingly, high parental mortality rates might not only increase the likelihood that these negative consequences arise; rather, they might also select against the further consolidation of family life, since the concomitant deepened integration of parental care into offspring development would increase the costs of parental loss. Similarly, the consolidation of family life might be hindered where (variation in) the availability of limited resources prevents parents from reliably provisioning their offspring. Such situations could also select for the maintenance of alternative survival strategies among juveniles (Kölliker, 2007; Kramer et al., 2015; Kramer & Meunier, 2016a). We surmise that the reliability of parental care – i.e. the likelihood that offspring indeed receive care once it has originated -will prove crucial in determining whether a given family system evolves towards increasing complexity (see also Capodeanu-Nägler et al., 2016).

Advanced family systems are typically caught in a *parental trap* that enforces the maintenance of family life irrespective of its current adaptive value (Eberhard, 1975). By contrast, less derived forms of family life can be lost over evolutionary times (Tallamy & Schaefer, 1997; Lin et al., 2004; Filippi et al., 2009). In the light of the above considerations, this contrast indicates the existence of a threshold of social complexity that determines whether family life is self-sustaining. Above this threshold, the phenotypic integration of parental care into offspring development would be tight enough to render parental care obligatory for offspring survival. Family life would then be beneficial to offspring irrespective of the external conditions, and could thus hardly ever be lost. By contrast, the integration of parental care into offspring development below this threshold would be sufficiently limited to enable offspring survival in the absence of the parents. In this situation, family life would remain facultative, and the interplay between environmental conditions, life-history characteristics and the costs-benefit ratio of all types of family interactions would determine whether family life is maintained at its *status quo*, abandoned in favour of a solitary lifestyle, or propelled towards the threshold that separates facultative from obligatory family systems. The existence of a similar threshold (or *point of no return)* has been invoked to explain the transition from facultative to obligatory eusociality (Wilson & Hölldobler, 2005). With regard to the evolution of family life, such a threshold would reconcile the current debate over the loss of parental care and family life (cf. Trumbo, 2012), since it allows for the co-existence of stable as well as unstable family systems. It would also leave scope for the theoretically expected unidirectional trend toward increasingly complex family systems – namely if the prevailing conditions are favourable and stable enough to promote an ever-increasing integration of parental and offspring traits.

## IV. IMPLICATIONS FOR SOCIAL EVOLUTION

Throughout the history of life on earth, previously independent units (such as cells) have formed social collectives (such as multicellular organisms) to cope with the challenges imposed by their changing environment. Transitions from solitary to social life were the incipient steps in such *major transitions* in evolution, and hence often had far-reaching repercussions on the diversity, complexity, and hierarchical organization of life itself (Maynard Smith & Szathmáry, 1995; Bourke, 2011). Indeed, the quest for general mechanism driving such transitions has prepossessed scientists ever since Charles R. Darwin (Darwin, 1859) first speculated on the evolution of eusocial societies (cf. Alexander, 1974; Krause & Ruxton, 2002; Bourke, 2011). Since then, the mechanisms driving transitions from simpler social systems to the highly integrated and often permanent societies of cooperatively breeding vertebrates and eusocial insects have been thoroughly explored (e.g. Wilson, 1971; Bourke & Franks, 1995; Crozier & Pamilo, 1996; Koenig & Dickinson, 2004, 2016). The evolutionary origin of the simpler social systems themselves, however, has received less attention (Trumbo, 2012; Falk et al., 2014; van Gestel & Tarnita, 2017; Boomsma & Gawne, 2017), and the mechanisms promoting the early evolution of social life remain poorly understood. The emergence of family living exemplifies a transition from solitary to social life, and marks the origin of an (initially) simple social system. Moreover, it constitutes the initial step towards the major transition to eusociality (Maynard Smith & Szathmáry, 1995; Bourke, 2011; Boomsma & Gawne, 2017). Understanding the origin and consolidation of family life might thus help to shed light on processes that also shape (the early steps of) other evolutionary transitions (see also van Gestel & Tarnita, 2017). In the following part, we discuss how adopting a broad perspective on the evolution of family life could provide general insights into the factors shaping social evolution.

### (1) Pathways to group formation

Social interactions among juveniles likely have a crucial impact on the early evolution of family units (see section III.1.b); yet their impact could go beyond the simple reinforcement of the benefits of parental care. In particular, the benefits of such interactions might influence the initial formation of family units, and could thus have implications for our understanding of the pathways to group formation. The transition to group-living is generally envisioned to follow either the semisocial or the subsocial pathway (Michener, 1969; Bourke, 2011). The semisocial pathway occurs when group formation results from the aggregation of individuals of the same generation, a process that, for instance, gave rise to the larval societies of sawflies and colonies of communally nesting halictid bees (Michener, 1969; Costa, 2006; Bourke, 2011). By contrast, the subsocial pathway occurs when group formation results from the association of parents with their offspring, an event that corresponds to the emergence of social interactions among the family members (Queller, 2000; Bourke, 2011), and ultimately gave rise to the majority of advanced animal societies (Wheeler, 1928; Wilson, 1975; Bourke, 2011; Boomsma & Gawne, 2017). Interestingly, the potential role of sibling cooperation during early stages of the evolution of family life (see section III.2.b) suggests that aggregations of juveniles might not only constitute an alternative (semisocial) pathway to group formation; rather, they could actually precede the emergence of (subsocial) family life. Specifically, semisocial aggregations of juveniles could initially arise whenever the benefits of sibling interactions favour delayed dispersal, and might subsequently give rise to families if parents extend already existing forms of pre-hatching care beyond offspring emergence (e.g. Lack, 1968; Clutton-Brock, 1991; Smiseth et al., 2012). This scenario suggests that species might not only exhibit both the subsocial and the semisocial pathway to group formation during different stages of their life cycle (Costa, 2006); rather, they might follow the two pathways at different times in the course of their evolutionary history.

### (2) The rise and fall of cooperation and conflict

In the course of major evolutionary transitions, cooperation typically spreads among lower-level units (such as individuals in the transition to eusociality) and replaces the initially prevailing conflicts between them (Bourke, 2011). The evolution of family life shows evidence for both processes: parental care, a hallmark cooperative trait (Hamilton, 1964; Smiseth et al., 2012), greatly diversifies during the evolution of complex family systems. Conversely, the initially prevailing direct competition between parents and offspring might be progressively suppressed (Kramer et al., 2017). However, the evolutionary dynamics shaping family living also indicate that not all forms of cooperation might be favoured and not all conflicts equally suppressed during its consolidation. For instance, cooperation among juvenile siblings might occur frequently in facultative family systems, but is arguably rare in advanced systems with obligatory family life (Roulin & Dreiss, 2012; Kramer et al., 2015). Conversely, sibling rivalry and parent-offspring conflict *(sensu* Trivers, 1974) typically increase during the evolution of complex family systems (Gardner & Smiseth, 2011). These findings suggest that some conflicts that are characteristic of later stages in an evolutionary transition might arise from dynamics that shaped earlier stages of that transition. In more general terms, they indicate that the increase in cooperation and the suppression of conflicts might be overall trends that need to hold true neither for all types of cooperation and conflict, nor for all stages of a transition. Notably, social systems might evolve towards a major transition even if a specific form of cooperation [such as sibling cooperation] is lost – namely if its benefits are offset by the benefits of a simultaneous increase in another form of cooperation [such as parental care] and/or the reduction in the costs of some form of conflict [such as parent-offspring competition].

### (3) The consolidation of social life

The various stages of a major transition broadly fall into two categories describing the initial formation of *collectives* (such as groups) out of formerly independent *particles* (such as individuals) on the one hand, and the subsequent transformation of these collectives on the other hand (Bourke, 2011). This transformational phase entails the transfer of key (e.g. metabolic or reproductive) functions from the particle to the collective level (Maynard Smith & Szathmáry, 1995; Bourke, 2011), and hence exhibits a striking resemblance to the consolidation of family life. In both cases, an increasingly tight phenotypic integration ties the fate of single particles [offspring] closer and closer to the fate of the collective [family], eventually resulting in obligatory social life – that is the inability of particles [offspring] to survive alone. This resemblance suggests that the reliability with which particles can derive benefits from the collective might have a crucial role in the transformational phase that corresponds to the role of the reliability of parental care in the consolidation of family life (see section III.2.c). For instance, the likelihood of a costly collapse of a facultative collective (i.e. the likelihood of ‘collective mortality’) might influence whether the phenotypic integration among its constituent particles proceeds, and could thus ultimately determine whether the collective becomes obligatory for particle-survival. Like the shift from facultative to obligatory family life, the shift from facultative to obligatory collectives could occur when environmental conditions and life-history characteristics of the particles allow for the breaching of a threshold of social complexity (see section III.2.b). Interestingly, the increasing phenotypic integration among the particles underlying this shift might also be paralleled by a shift from particle to collective-level selection (Okasha, 2005; Shelton & Michod, 2010). This change in the most relevant level of selection could in turn determine whether kin selection or multilevel selection approaches best describe the underlying evolutionary process (Kramer & Meunier, 2016b; Okasha, 2016). The different stages of the evolution of family life offer rich opportunities to investigate these possibilities. Exploring the intricacies of family life might thus be a good starting point to advance our understanding of the major transitions and the theoretical framework of sociobiology.

## VI. CONCLUSIONS

(1) Over the last decades, the intricacies of family interactions received theoretical and empirical scrutiny in a plethora of studies that focused on parental care and its associated family interactions (such as those arising from sibling rivalry and parent-offspring conflict), and investigated these phenomena in altricial vertebrates and eusocial insects. This historical bias bears on the often-substantial fitness effects of these phenomena in derived family systems. However, it has led to a neglect of mechanisms that might be particularly important in shaping the social life in less-derived family systems. Consequently, a coherent framework for the study of social interactions and fitness effects of family life is currently missing, and our understanding of the (early) evolution of family life remains limited.

(2) Here, we argued that the explicit consideration of thus far neglected facets of family life – and their study across the whole taxonomical diversity of family systems – is crucial to shed light on the mechanisms driving the evolution of social life in family groups. In particular, we illustrated that the strong focus on parental care in advanced social systems has fostered the neglect of three facets of family life: sibling cooperation, parent-offspring competition, and offspring assistance. We suggested that the impact of these facets is often – and especially in derived family systems – concealed by the fitness effects of parental care.

(3) We showed how accounting for these overlooked facets – and their changing role in the course of evolution – is nevertheless crucial, and could improve our understanding of the evolutionary emergence and consolidation of family life. Specifically, we highlight that both sibling cooperation and offspring assistance could promote the evolutionary emergence of family life by, respectively, augmenting the benefits and offsetting some of the costs of parental care. Conversely, we suggest that parent-offspring competition might impede the evolution of family life by reducing the net benefits of care. We argue that all three thus far neglected facets have a greater impact where offspring are largely independent of (and thus do not compete for) parental care – a scenario that prevailed during the early evolution of family life, and is prevalent among contemporary precocial species and in adult offspring of cooperative breeders.

(4) We show that the study of family interactions in (precocial) species featuring non-derived forms of family life is not restricted to elucidating the role of sibling cooperation, parent-offspring competition, and offspring assistance; rather it can also shed light on factors – such as the reliability of the benefits of parental care – that can affect the benefits of a (further) consolidation of family life, and thus promote or hamper the evolution of complex animal societies.

(5) Finally, we discuss how diachronic perspective on the evolution of family living could provide novel insights into the mechanisms driving social evolution. In particular, we suggest that (subsocial) family life can evolve secondarily from semisocial aggregations of juveniles that delay dispersal to reap the benefits of sibling cooperation. We argue that the role of the reliability of the benefits of parental care in the consolidation of family life can be generalized, which would suggest a key role of the reliability of ‘collective’ benefits in the consolidation of social life.

(6) Overall, we aimed at providing a general perspective on the evolution of family life that accounts for all types of family interaction across the whole taxonomical diversity of family systems. Recent advances in the study of parental care stress its multifaceted nature (e.g. Gardner & Smiseth, 2011; Royle et al., 2016; Andrews, Kruuk, & Smiseth, 2017); we hope that our perspective on the intricacies of family life complements this fruitful trend by raising awareness for the multifaceted nature of social life in family groups. The further development of this perspective hinges on studies that investigate family life in species with non-derived (facultative) forms of family life. Many allegedly ‘primitively social’ insects (see Tallamy & Wood, 1986; Costa, 2006; Trumbo, 2012; Wong et al., 2013 for reviews) offer unprecedented opportunities to study the origin and maintenance of early forms of parental care and family life (Smiseth et al. 2003b; Kölliker 2007; Trumbo 2012). We believe that their resemblance to ancestral family systems, and the great diversity of family interactions across species, could well render them prime models of social evolution.

## VII. ACKNOWLEDGEMENTS

The authors thank Maximilian Körner for his comments on a previous version of this manuscript. JK was supported by a grant from the University of Zürich (Forschungskredit of the University of Zurich, grant no. [FK-17-111]).

